# Intergenerational transmission of the patterns of functional and structural brain networks

**DOI:** 10.1101/2020.03.06.981068

**Authors:** Yu Takagi, Naohiro Okada, Shuntaro Ando, Noriaki Yahata, Kentaro Morita, Daisuke Koshiyama, Shintaro Kawakami, Kingo Sawada, Shinsuke Koike, Kaori Endo, Syudo Yamasaki, Atsushi Nishida, Kiyoto Kasai, Saori C Tanaka

## Abstract

There is clear evidence of intergenerational transmission of life values, cognitive traits, psychiatric disorders, and even aspects of daily decision making. To investigate biological substrates of this phenomenon, brain has received increasing attention as a measurable biomarker and potential target for intervention. However, no previous study has quantitatively and comprehensively investigated the effects of intergenerational transmission on functional and structural brain networks from parents to their children. Here, by employing an unusually large cohort dataset, we show that patterns of functional and structural brain networks are preserved over a generation. We also demonstrate that several demographic and behavioural phenotypes have effects on brain similarity. Collectively, our results provide a comprehensive picture of neurobiological substrates of parent-child similarity, and demonstrate the usability of our dataset for investigating the neurobiological substrates of intergenerational transmission.

## Introduction

There is clear evidence of intergenerational transmission of socio-economic status^1^, intelligence^2^, personality^3^, parenting style^4^, job-selection^5^, and psychiatric disorders^6^. This correspondence between parents and their children is not confined to the period in which children are young and live with their parents, but is found over the course of their lives^7^. Although genetic and non-genetic environmental effects are clearly transferred to children from their parents, the mechanisms of parent-child similarity are poorly understood^8^.

In recent years, brain has received increasing attention as a target for monitoring and intervention because genetic and epigenetic effects occur at the molecular level and are distal from complex behaviour^9^. Several previous studies have shown that functional connectivity (FC or edge) during wakeful rest obtained by functional magnetic resonance imaging (fMRI) is associated with individual differences in diverse cognitive traits^10–19^. In parallel to functional brain information, individual differences in brain structure have also been characterised and related to diverse cognitive traits^20^. Importantly, previous studies have reported that grey matter volume (GMV) at specific locations in the brain is associated with individual differences in cognitive traits^20–24^.

In addition to individual differences in cognitive traits, previous studies also showed that FC^25–43^ and GMV^25, 44–57^ are heritable. These studies have typically used genome wide association study (GWAS) or family/twin study. Most studies have assessed the genetic effects on each edge- or region-level, i.e. univariate analysis, and typically considered demographic or behavioural information as covariates. Importantly, previous studies have not directly focused on the effects of intergenerational transmission from parents to their children.

More recent studies have started to directly examine the effects of intergenerational transmission on the brain using datasets of parent-child dyads^8, 59–62^. For example, Lee et al. and Yamagata et al. investigated the similarity of parent-child dyads in FC and GMV, respectively^59,62^. However, these studies involved several limitations. First, no study has quantitatively compared the similarity of different brain networks in detail. Second, because none of these studies examined both functional and structural data together, it remains unclear how functional and structural information are interrelated. Third, no study has comprehensively investigated the effects of demographic and behavioural effects on similarity. Overall, it is currently unclear whether, to what extent, and how the brains of parent-child dyads are similar. This situation has arisen, in part, because investigating the above questions requires a large number of parent-child dyads to provide neuroimaging datasets with rich behavioural phenotypes. Furthermore, such an approach requires a formal analytical framework with rigorous statistical analyses and rich computational resources.

In the current study, we sought to understand the neurobiological substrates of parent-child similarity by combining a statistical framework that allowed us to investigate network-level similarities and a rich dataset from a subsample of a large population-based longitudinal cohort (N = 84 parent-child dyads) consisting of resting-state fMRI, structural MRI, and behavioural phenotypes of parents and their children^63,64^. We sought to answer several questions: Can a parent-child dyad be identified based on their brains? If so, which brain networks are more similar compared with other networks? Is the similarity solely driven by functional or structural information? How do demographic and behavioural factors affect brain similarity?

Using a dataset consisting of parents and their children, we quantitatively investigated the brain similarity of parent-child dyads in detail. The present findings demonstrated that it is possible to reliably identify a parent-child dyad based on the similarity of their brains. This effect was not solely driven by either functional or structural brain similarity alone: although functional and structural information had comparable accuracy, they exhibited important differences, and played complementary roles. Children’s basic demographic factors, including age and sex, testosterone level, and questionnaire-based developmental scores affected parent-child brain similarity. Collectively, our results provide a detailed picture of how the brains of children and their parents are similar, and demonstrate the usability of our unique cohort dataset for investigating the neurobiological substrates of intergenerational transmission.

## Results

We tested 84 parent-child dyads who participated in the “population-neuroscience study of the Tokyo TEEN Cohort (pn-TTC),” a longitudinal study exploring the neurobiological substrates of development during adolescence^63,64^. In the pn-TTC study, neuroimaging and non-imaging behavioural phenotypes were collected from children every 2 years from the age of 11, and from their primary parents (Figure 1a; see *Methods: Overview of the dataset*). Here, we used three brain datasets from the pn-TTC study: children at the ages of 11 and 13 years, and their primary parents. Parents’ brains were scanned when their children were 11 years old. The basic demographic data are shown in Table 1.

**Figure 1:**
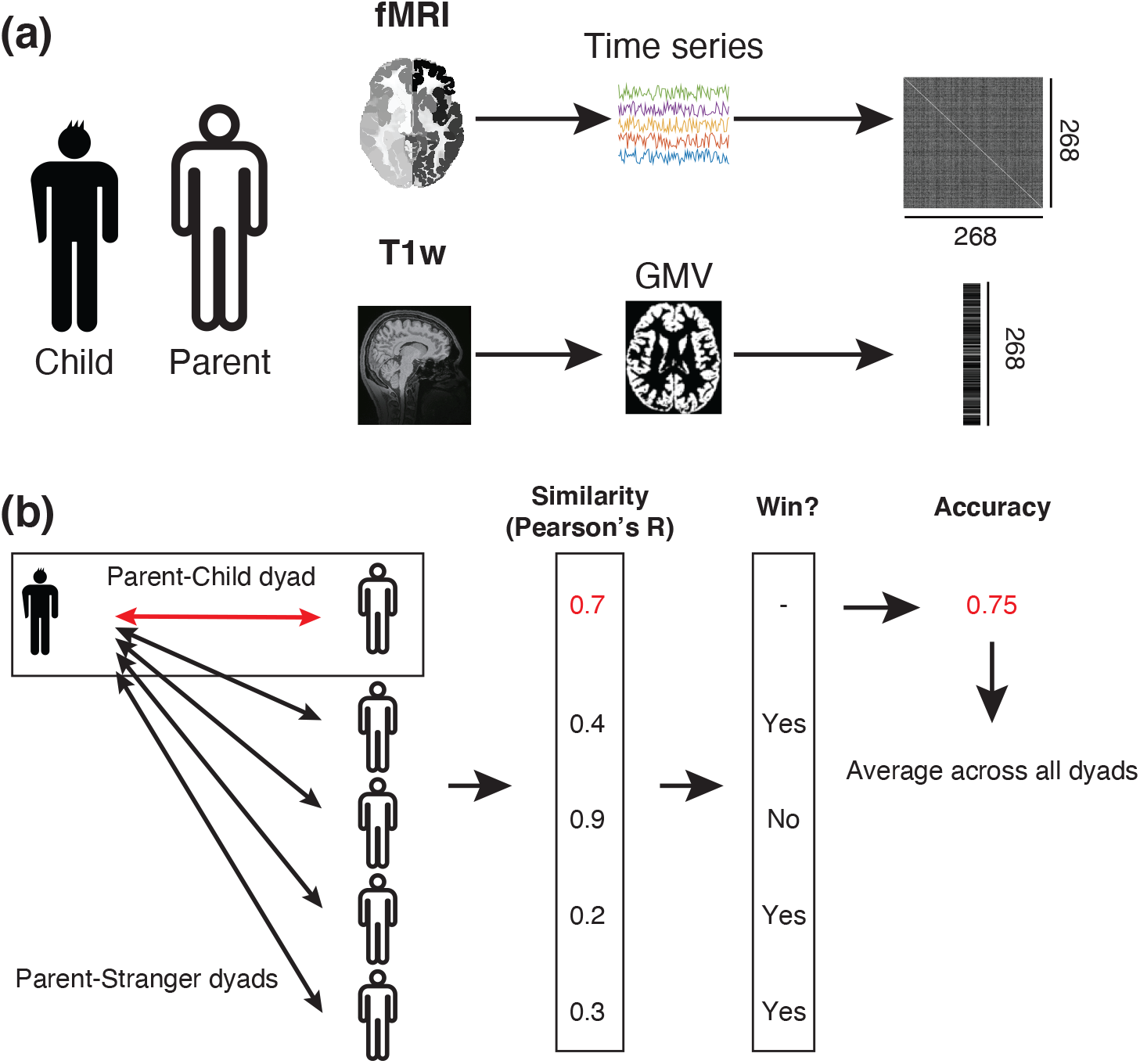
Analysis procedure of parent-child brain similarity. (**a**) We employed a dataset obtained from the “population-neuroscience study of the Tokyo TEEN Cohort (pn-TTC)” study, which consists of resting-state fMRI and T1w images of parents and their children. To obtain a functional connectivity (FC) matrix, signals were extracted from all subjects using resting-state fMRI data from 268 ROIs. The signals were then turned into an FC matrix via covariance estimation. To obtain grey matter volume (GMV) vectors, T1w images were first segmented into grey matter, white matter, and cerebrospinal fluid. Using the grey matter, GMV of each ROI was obtained by averaging values within the ROI. We used the same 268 ROIs as in the FC. **(b)** For each parent-child dyad, we first calculated the similarity of FC and/or GMV vectors based on their Pearson’s correlation. We next calculated similarities between the child and another child’s parent. We then calculated whether the similarity of the parent-child dyad is greater than that of stranger-child dyads. Finally, we calculated the winning rate of the parent-child dyad (“accuracy”), and repeated this procedure across all parent-child dyads.

**Table 1:** Demographic data. ^a^ Ages of parents when their children were 11 years old are shown.

| | N | Age<sup>a</sup><br>(mean ± s.t.d) | Sex<br>(Male/Female) |
| --- | --- | --- | --- |
| Child age at 11 | 84 | 11.59±0.66 | 45/39 |
| Child age at 13 | | 13.63±0.62 | |
| Parent | | 43.35±3.95 | 3/81 |

We first defined the functional and structural whole-brain patterns for each individual (Figure 1a; see *Methods: Information extraction*). For fMRI, we used a functional atlas defining 268 regions of interest (ROIs) covering the entire brain^12,65^. The FC between these ROIs was estimated using Pearson’s correlation coefficient, resulting in a 268 × 268 FC matrix for each subject. As the FC matrix is symmetrical, only the strictly lower triangular part of each matrix was kept, resulting in 35,778 (= 268 × 267 / 2) unique entries. We regressed potential confounds including total GMV and head motion. To further avoid the effects of motion artefacts, we employed a “scrubbing” procedure to identify and exclude any frames exhibiting excessive head motion^66^. We also obtained GMV for each ROI using T1w images, then averaged within each region. We used the same 268 ROIs as in the fMRI, resulting in a vector with a size of 268 for each subject.

To quantitatively evaluate the brain similarity of parent-child dyads, we proposed a framework to compare network-level similarity between parent-child dyads (Figure 1b; See *Methods: Similarity analysis* for details), inspired by a recently proposed approach for individual identification based on the brain^12^. To calculate the similarity of parent-child dyads, we first calculated the correlations between a child’s FC and/or GMV vector to all parents’ vectors including the child’s own parent. We next assessed whether the similarity of the parent-child dyad (child and his/her own parent) was larger than that of a stranger-child dyad (child and another child’s parent). We then calculated the winning rate of the similarity between parent-child dyad, which was referred to as “accuracy”, because it can be considered as a “pairwise classification accuracy” when we randomly sampled a parent-child dyad and another parent, then conduct classification (See *Methods: Similarity analysis* for details). We repeated this procedure across all dyads and averaged these accuracies. Compared with conventional individual identification methods, our proposed framework has more statistical power, as described later. We performed 1,000-times bootstrapping to estimate 95% confidence intervals of accuracy by randomly subsampling 90% of the subjects in each iteration. To determine whether accuracy was achieved at above-chance levels, we used 1,000-times permutation testing to generate a null distribution by randomly shuffling the parent-child mapping.

### Whole-brain analysis

We first assessed the similarity of parent-child dyads using whole-brain FC and GMV. When we used a dataset of children at age 11, accuracies were 64.6% for FC (estimated via 1,000-times bootstrapping; 95% CI = [62.5, 66.8]; P < 0.001, 1,000-times permutation test) and 70.3% for GMV (95% CI = [68.9, 71.8]; P < 0.001) (Figure 2a). When we used a dataset of children at age 13, the accuracies were 66.7% for FC (95% CI = [64.5, 68.9]; P < 0.001) and 73.8% for GMV (95% CI = [72.1, 75.6]; P < 0.001) (Figure 2b). Thus, we provided the first strong evidence that it is possible to identify parent-child dyads based on their functional and structural brain information.

**Figure 2:**
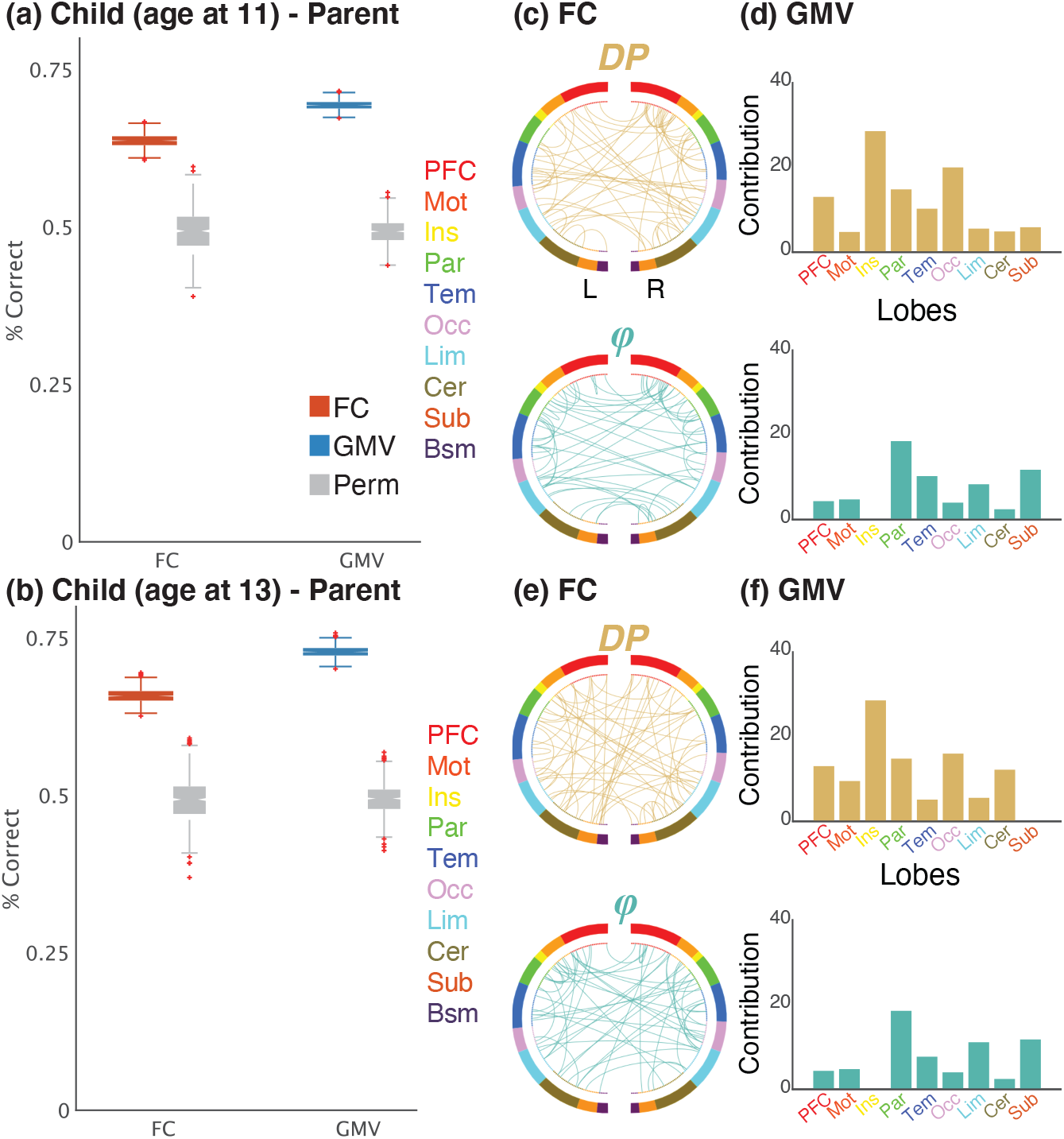
Successful identification of parent-child dyad based on their functional and structural brain information. Box plots of parent-child identification accuracy and factors affecting accuracy for **(a)** children at age 11 and **(b)** at age 13 using whole-brain (268-node) for functional connectivity (FC: red box) and grey matter volume (GMV: blue box). Directly to the right of these boxes (grey box) are the results of the 1,000-times permutation testing. The bottom and top edges of the box indicate the 25th and 75th percentiles obtained via bootstrapping, respectively. The crosses denote outliers, and the whiskers extend to the most extreme data points not considered outliers. **(c–f)** Factors affecting identification accuracy. For FC, the top 99.75th percentile of differential power (DP: highly discriminative; yellow) edges and group consistency (*φ*: highly similar, or least helpful; green) edges are shown (circle plot; in which nodes are grouped according to anatomic location). For GMV, the top 90th percentile ROIs of DP and *φ* were calculated, then normalised by dividing the number of ROIs in each anatomical group (bar plot). A legend indicating the approximate anatomical “lobe” is shown. PFC, prefrontal; Mot, motor; Ins, insula; Par, parietal; Tem, temporal; Occ, occipital; Lim, limbic (including cingulate cortex, amygdala, and hippocampus); Cer, cerebellum; Sub, subcortical (including thalamus and striatum); Bsm, brainstem; L, left hemisphere; R, right hemisphere.

We next assessed the importance of information for performance of specific edges for FC and regions for GMV, respectively (Figures 2c-2f). To quantify the extent to which different edges and regions contribute to similarity, we derived two measures: the differential power (DP) which calculates how characteristic edges and regions tend to be, and group consistency (*φ*) which quantifies edges and regions that are highly consistent across all parent-child pairs in a dataset^12^ (see *Methods: Similarity analysis*). For visualisation purposes, we show the structural locations of DP and *φ* in the 99.75th and 90th percentile, for edges (FC) and regions (GMV) respectively. For both FC and GMV, significant edges or regions tended to be distributed across the entire brain. This pattern was stable across a range of thresholds (Supplementary Figure 1). Note that, for visualisation purposes, we excluded the brainstem from the figure for GMV because all regions in this area had extremely high *φ* values, possibly because of the much lower magnitudes of signals compared with the other regions (Supplementary Figure 2).

Given that head motion confounds analyses of FC^66^, we confirmed that qualitatively similar results were obtained when we excluded parent-child dyads whose children’s head movements were in the top 25%, either at age 11 or 13, resulting in the inclusion of 41.25% of the total sample (Supplementary Figure 3). We also confirmed that accuracy obtained by the distribution of their frame-to-frame motion during fMRI scans^12^ was not above chance level (51.0% for age 11, 95% CI = [48.9, 53.0]; 48.7% for age at 13, 95% CI = [47.1, 50.1]). Thus, it is unlikely that the identification power of FC was based on idiosyncratic patterns related to motion in the scanner.

### Network-based similarity

We further investigated the contributions of specific networks to this similarity. We grouped the whole regions into 10 sub-networks^67^ (Figure 3a), and subsequently performed the same analyses using only the edges and regions from a given network. Note that we calculated the null distribution for each network via permutation testing, thus taking differences of the number of edges/regions among networks into account.

**Figure 3:**
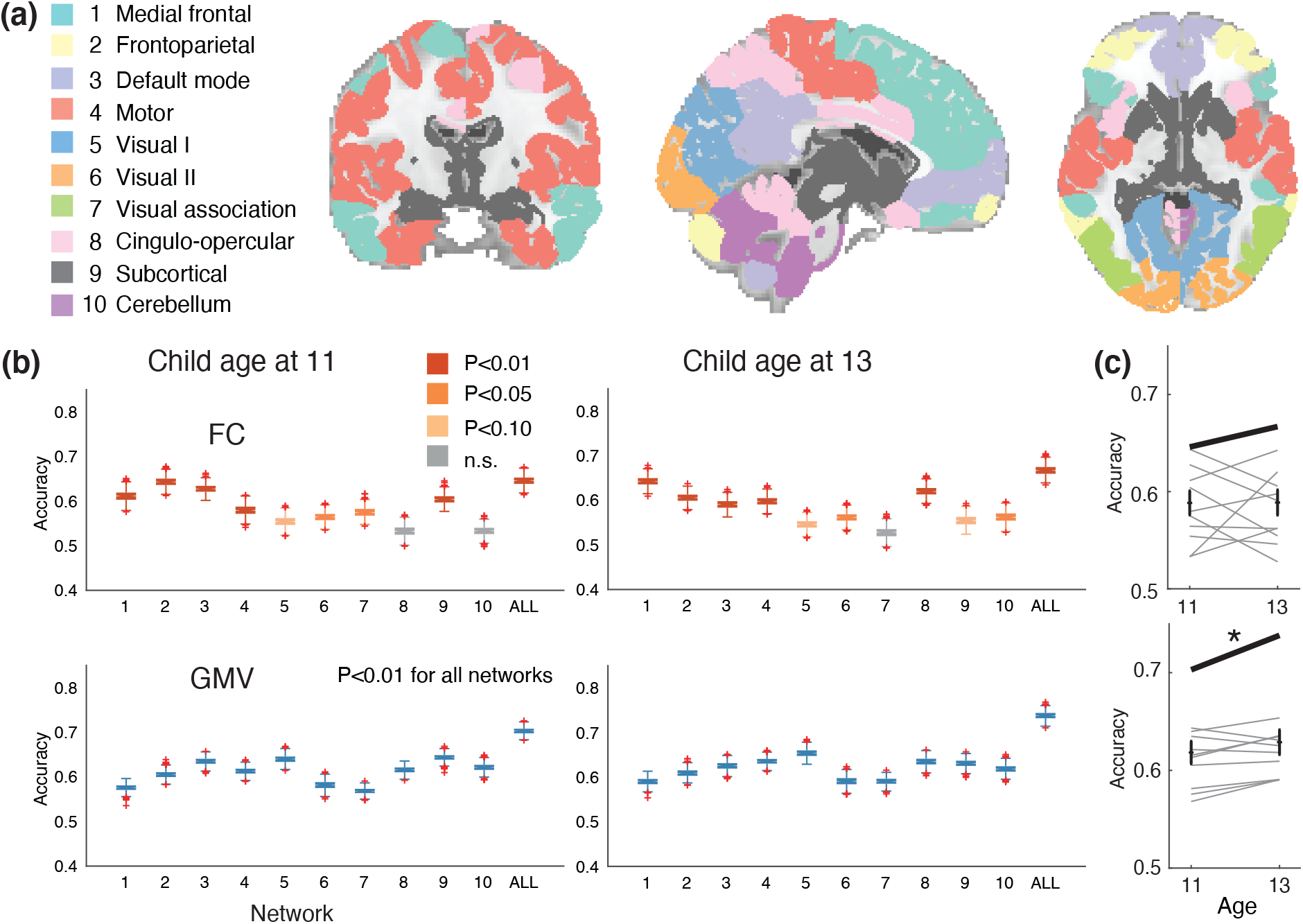
Network-based analyses demonstrated that almost all brain networks were highly similar between parents and their children. (**a**) We utilised a 268-node functional atlas. Nodes were further grouped into the 10 functional networks. Network names are shown to the left. (**b**) Box plots of accuracies using within-network edges of FC analysis (top row; networks 1–10 and whole-brain (ALL); indicated below the x-axis of each graph) and within-network nodes of GMV (bottom row). The bottom and top edges of the box indicate the 25th and 75th percentiles, respectively. The crosses denote outliers, and the whiskers extend to the most extreme data points not considered outliers. (**c**) Comparison between accuracies of child age at 11 and 13 for FC (top row) and GMV (bottom row). Each scatter shows each network and line connected the same network. Bold lines indicate ALL. * paired sample t-test, P < 0.05, n.s. non-significant, uncorrected.

For FC, medial frontal and frontoparietal networks led to high accuracies (Figure 3b). In contrast, for GMV, default mode, subcortical, cerebellum and visual networks led to high accuracy. Compared with FC, GMV achieved modestly higher accuracy than FC at age 11 (Figure 3c; paired sample t-test, t(10) = −2.16, P = 0.056, Hedge’s g = −0.72) and significantly higher accuracy at age 13 (Figure 3c; paired sample t-test, t(10) = −3.10, P = 0.011, Hedge’s g = −0.89). To further assess the importance of each network, we next assessed performance using between-network pairs of edges. We observed that edges between the medial frontal– frontoparietal networks and medial frontal–motor networks resulted in higher accuracies than the other between-network pairs (Supplementary Figure 4).

The results confirmed that our proposed method had greater statistical power than conventional methods for individual identification (Supplementary Figure 5). Specifically, for FC, six and five of 11 networks were significant at age 11 and 13, respectively, using the conventional method, whereas nine and 10 networks, respectively, were significant in our proposed method. For GMV, six and nine of 11 networks were significant at age 11 and 13, respectively, using the conventional method, whereas all networks were significant using our proposed method.

### Function and structure provide complementary information

Although FC and GMV revealed comparable performance in the above analyses, it remained unclear whether they contained similar information. This raises the following question: if parents and their children exhibit similar patterns of structural brain information, do they also exhibit similar patterns of functional brain information? Indeed, although functional and structural brain information is interrelated, they contain exclusive information that putatively characterises distinct properties of individual differences^68^. To test this question, we investigated the relationship between parent-child similarity defined by FC and that of GMV. We found that the Pearson’s correlation between the similarities defined by FC and GMV was low: only two of 22 networks were significant (Supplementary Figure 6; P = 0.026 for visual and P = 0.021 for whole-brain when we used data from children at age 13; the other networks were not significant, P > 0.05, 1,000-times permutation test, uncorrected). Thus, although both FC and GMV were similar between parent-child dyads, their characteristics were dissimilar.

Given that FC and GMV appeared to contain independent information, we further investigated whether they contained complementary information. To test this question, we conducted the same analyses using both FC and GMV simultaneously by concatenating the two vectors (hereafter referred to as “COMB”). COMB achieved the highest accuracies in more than half of cases (15/22), compared with function (4/22) and structure (3/22) alone (Table 2). This number is significantly greater than chance (Supplementary Figure 7; P < 0.001, 1,000-times permutation test). Overall, these results suggest that patterns of functional and structural information contained complementary information in terms of parent-child brain similarity.

**Table 2:** COMB achieved higher accuracies compared with FC and GMV for many networks in children at age 11 and at age 13. Means and 95% confidence intervals estimated via 1,000-times bootstrapping are shown. **<u>Bold underlined</u>** text indicates the best performance. MF, medial frontal; FP, frontoparietal; DMN, default mode network; Mot, motor; Vis I, visual I; Vis II, visual II; Vis A, visual association; Cing, cingulo-opercular; Sub, subcortical; Cer, cerebellum.

| Network | Age | FC | GMV | COMB |
| --- | --- | --- | --- | --- |
| ALL | 11 | 64.60 [62.54,66.83] | 70.30 [68.86,71.75] | **<u>70.64 [69.21,72.16]</u>** |
|  | 13 | 66.69 [64.50,68.86] | **<u>73.82 [72.14,75.55]</u>** | 73.57 [71.89,75.30] |
| 1. MF | 11 | **<u>61.14 [58.83,63.66]</u>** | 57.56 [56.02,58.94] | 58.32 [56.77,59.66] |
|  | 13 | **<u>64.26 [62.20,66.36]</u>** | 59.02 [57.06,60.70] | 59.14 [57.14,60.92] |
| 2. FP | 11 | **<u>64.35 [62.40,66.52]</u>** | 60.52 [59.01,62.20] | 61.90 [60.38,63.55] |
|  | 13 | 60.57 [58.61,62.49] | 60.94 [59.35,62.65] | **<u>61.79 [60.25,63.46]</u>** |
| 3. DMN | 11 | 62.77 [61.06,64.65] | 63.49 [61.93,64.94] | **<u>63.66 [62.04,65.21]</u>** |
|  | 13 | 59.07 [57.03,61.03] | 62.53 [60.77,64.23] | **<u>62.84 [61.05,64.50]</u>** |
| 4. Mot | 11 | 57.95 [55.60,60.29] | 61.30 [59.80,62.85] | **<u>62.27 [60.70,63.82]</u>** |
|  | 13 | 59.78 [57.89,61.77] | 63.56 [61.95,65.14] | **<u>64.27 [62.63,65.98]</u>** |
| 5. Vis I | 11 | 55.42 [53.19,57.59] | 63.98 [62.40,65.59] | **<u>64.91 [63.23,66.58]</u>** |
|  | 13 | 54.60 [52.61,56.77] | 65.37 [63.60,67.21] | **<u>65.98 [64.20,67.77]</u>** |
| 6. Vis II | 11 | 56.44 [54.41,58.61] | 58.14 [56.20,60.04] | **<u>58.95 [57.14,60.81]</u>** |
|  | 13 | 56.20 [54.14,58.25] | 59.07 [56.95,61.03] | **<u>60.33 [58.22,62.32]</u>** |
| 7. Vis A | 11 | **<u>57.52 [55.35,59.86]</u>** | 56.81 [55.51,58.02] | 57.13 [55.84,58.31] |
|  | 13 | 52.78 [50.50,55.06] | **<u>59.05 [57.28,60.56]</u>** | 58.92 [57.28,60.41] |
| 8. Cing | 11 | 53.28 [51.08,55.66] | **<u>61.54 [59.89,62.99]</u>** | 61.53 [59.91,62.94] |
|  | 13 | 62.04 [59.93,64.34] | 63.48 [61.51,65.17] | **<u>63.71 [61.68,65.46]</u>** |
| 9. Sub | 11 | 60.43 [58.52,62.59] | 64.34 [62.67,65.95] | **<u>64.66 [62.97,66.23]</u>** |
|  | 13 | 55.45 [53.23,57.62] | 63.08 [61.41,64.65] | **<u>63.61 [62.00,65.19]</u>** |
| 10. Cer | 11 | 53.32 [51.23,55.37] | 62.13 [60.38,63.73] | **<u>62.86 [61.24,64.54]</u>** |
|  | 13 | 56.24 [53.87,58.68] | 61.83 [59.91,63.71] | **<u>62.50 [60.67,64.34]</u>** |
| # best |  | 4 | 3 | 15 |

### Effects of demographic factors on the brain similarity

The results described above indicated strong intergenerational transmission effects on the brain from parents to their children. However, it was still unclear whether all parent-child dyads were equally similar. Therefore, we then investigated which factors influence brain similarity. Here, we focused on fundamental demographic factors: age and sex.

We found that accuracies at age 11 were significantly lower than those at age 13 for GMV (Figure 3c bottom; paired sample t-test, t(10) = −2.39, P = 0.038, Hedge’s g = −0.25), whereas they were no different for FC (Figure 3c top; paired sample t-test, t(10) = −0.03, P = 0.97, Hedge’s g = −0.01).

We then divided children into males and females (Figure 4). The results confirmed that both male and female children exhibited significant accuracies for almost all networks, as in the previous analyses (Figure 4a). When we compared males and females (Figure 4b), the accuracies of female children were significantly higher than those of male children for FC, at both age 11 (paired sample t-test, t(10) = −2.34, P = 0.04, Hedge’s g = −0.88) and age 13 (paired sample t-test, t(10) = −3.80, P = 0.003, Hedge’s g = −1.21). Female children also exhibited higher accuracy than male children for GMV at age 13 (paired sample t-test, t(10) = −2.47, P = 0.03, Hedge’s g = −0.83), but not at age 11 (paired sample t-test, t(10) = −0.25, P = 0.81, Hedge’s g = 0.07).

**Figure 4:**
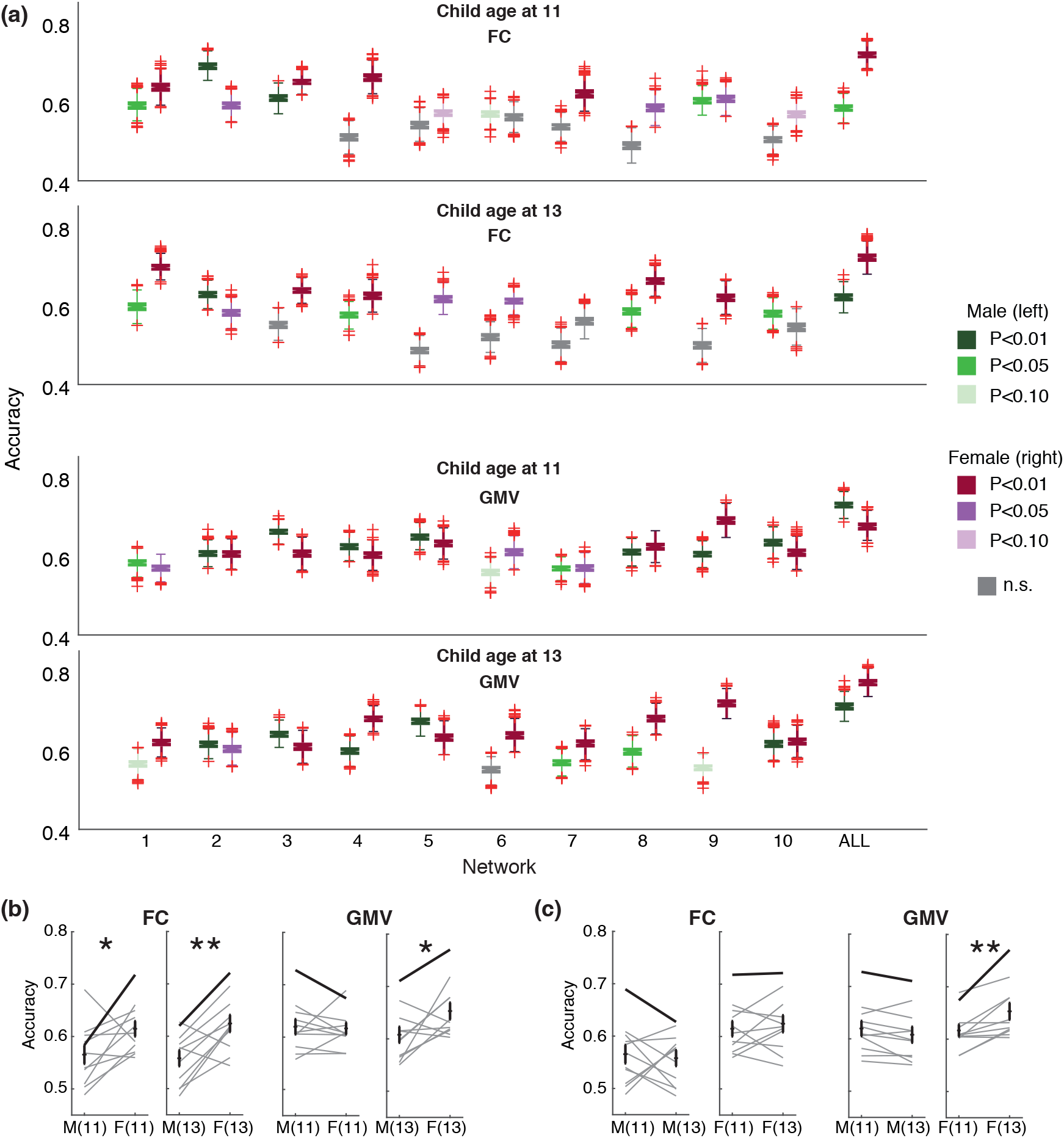
Male and female children show different trends of brain similarity across development. (**a**) Accuracies split by children’s sex for FC and GMV. For each network, accuracies for males (left boxes) and females (right boxes) are shown. (**b**) Comparison between males and females. (**c**) Comparison between children age at 11 and 13. Each scatter shows each network and line connected the same network. Bold lines indicate ALL. M(11), males at age 11; F(11), females at age 11; M(13), males at age 13; M(13), males at age 13; * Paired sample t-test, P < 0.05; ** P < 0.01. n.s. non-significant, uncorrected.

When we compared accuracies at age 11 and age 13 for male and female children separately (Figure 4c), female children at age 13 had significantly greater accuracy than those at age 11 for GMV (paired sample t-test, t(10) = −3.75, P = 0.004, Hedge’s g = −0.76). All other comparisons were not significant: male, FC (paired sample t-test, t(10) = 0.37, P = 0.72, Hedge’s g = 0.13); male, GMV (paired sample t-test, t(10) = 2.05, P = 0.068, Hedge’s g = 0.23); female, FC (paired sample t-test, t(10) = −0.73, P = 0.48, Hedge’s g = −0.18)

### Effects of behavioural phenotypes on brain similarity

Finally, we examined whether behavioural phenotypes have effects on the brain similarity of parent-child dyads. Here, we used two important behavioural phenotypes for adolescents: hormone level and questionnaire-based developmental score (Figure 5; see *Methods: Testosterone and Child Behavior Checklist*). We used COMB for this analysis as brain information.

**Figure 5:**
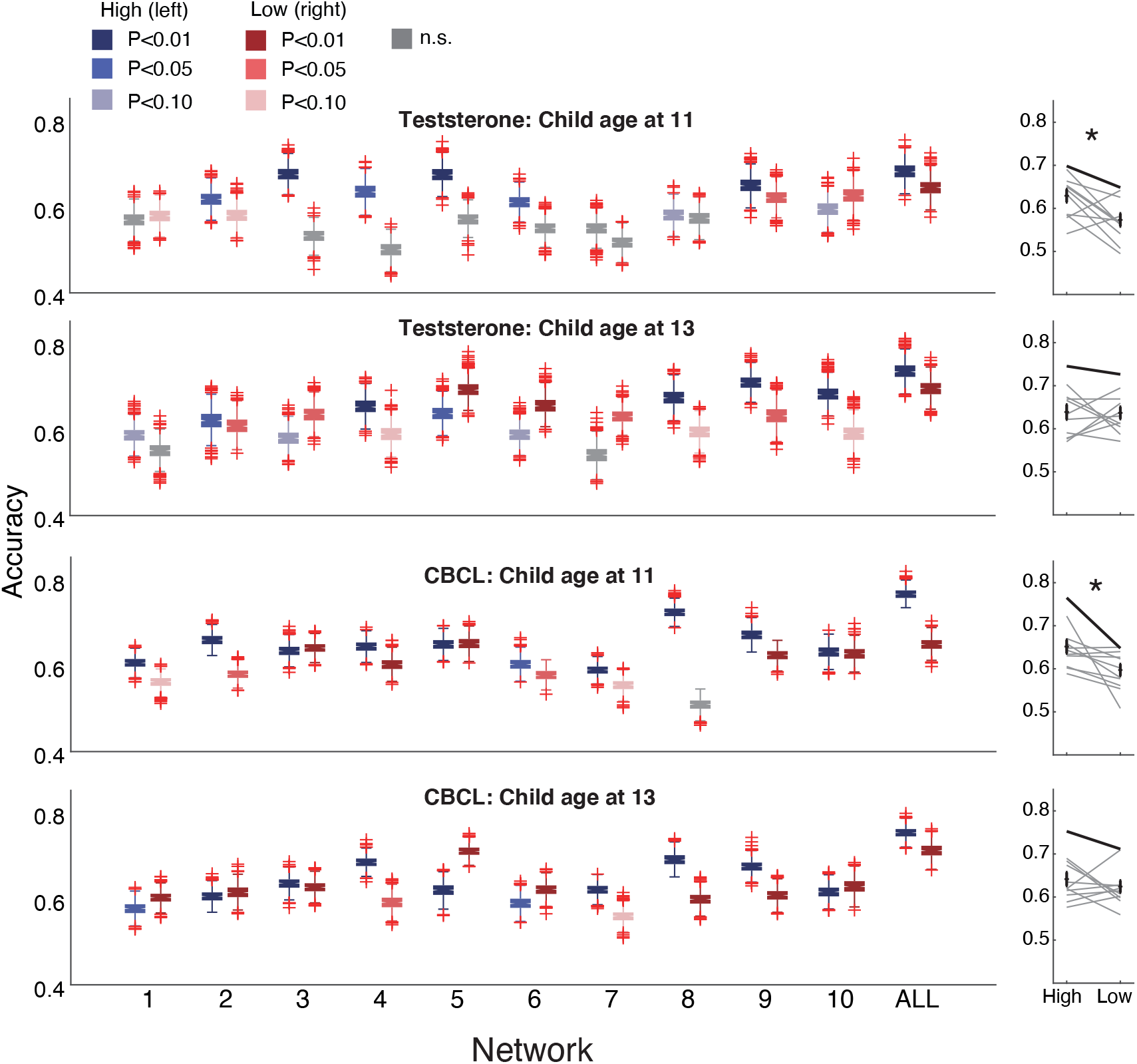
Behavioural phenotypes have effects on brain similarity. Accuracies split by testosterone and CBCL. Left panels: Boxplots of accuracies split by scores of behavioural phenotypes (left: upper-half children; right: lower-half children). Right panels: Comparisons between upper- (left) and lower-half (right) children. Each scatter shows each network and line connected the same network. Bold lines indicate ALL. * Paired sample t-test, P < 0.05. n.s. non-significant, uncorrected.

We examined testosterone as a hormone level, because it is known that the pubertal period is a sensitive period for testosterone-dependent organisation of the brain^69^. We found that children with high testosterone exhibited significantly higher accuracy compared with children with low testosterone at age 11 (paired sample t-test, t(10) = −2.91, P = 0.016, Hedge’s g = −1.03) but not at age 13 (paired sample t-test, t(10) = −0.60, P = 0.56, Hedge’s g = −0.22).

We next investigated the effects of a questionnaire-based development score (Child Behavior Checklist: CBCL)^70^. The CBCL is a parental-report assessment used to screen for emotional, behavioural, and social problems, and to predict psychiatric illnesses^71^. We found that children with high CBCL had significantly higher accuracy compared with children with low CBCL at age 11 (paired sample t-test, t(10) = −2.81, P = 0.018, Hedge’s g = −1.05) but not at age 13 (paired sample t-test, t(10) = −0.99, P = 0.34, Hedge’s g =−0.33).

## Discussion

The present study revealed that patterns of functional and structural brain information are preserved over a generation. Although the effects of the intergenerational transmission have been investigated in various research fields, including developmental psychology, educational psychology, and economics, no study has comprehensively and quantitatively investigated the neurobiological substrates of intergenerational transmission. The current results revealed that, despite substantial differences between parents and their children, their brains are sufficiently similar that we can identify parent-child dyads based on information about their brains. We employed a rigorous statistical framework and an unusually large functional and structural neuroimaging dataset of parent-child dyads with rich behavioural phenotypes (N = 84 parent-child dyads)^63,64^. Although both functional and structural brain information has comparable levels of accuracy, their characteristics were different but complementary. Demographic factors and behavioural phenotypes also have large effects on brain similarity. Taken together, our results provide a detailed picture of whether, to what extent, and how brains of parent-child dyads are similar.

Previous studies have reported that brain information is heritable. Several genome-wide association studies (GWAS) have been conducted to identify genetic risk variants for GMV^25,44,50–56^ and FC^25,26^. Although heritable regions or edges have been successfully identified, most studies have been insufficiently powered because GWAS require a large sample size (although some studies used multivariate approaches to increase statistical power^26,58^). In addition, these studies typically did not consider the effects of demographic and behavioural phenotypes. In addition to GWAS, both twin and family-based studies have reported that GMV^45–49,57^ and FC^27–43^ are heritable. These studies typically achieve larger effect sizes than GWAS studies with smaller sample sizes, although the possibility of inflated effect sizes due to shared environments is a concern. These studies also typically ignore demographic and behavioural phenotypes by treating them as covariates. Importantly, none of the previous studies described here directly investigated the effects of intergenerational transmission from parents to children.

In recent years, some studies directly investigated intergenerational transmission of brain information using parent-child dyads. Although extended pedigree studies with sufficient sample sizes could answer such a question, it is logistically more difficult to recruit participants for pedigree studies than for studies with a parent-child design. Thus, parent-child design studies play a complementary role in the investigation of intergenerational transmission. For example, some studies found that, during specific tasks, brain activity of parent-child dyads was synchronised^60,61^. Other, more relevant studies to the current studies reported that patterns of FC ^59^ and GMV ^62^ were similar between parent-child dyads. However, these studies did not quantitatively and comprehensively investigate the effects of different brain networks, and did not compare FC and GMV. It is also unclear which factors affect brain similarities, including age, sex, hormonal level, and behavioural traits (although Yamagata et al. investigated the effects of sex on GMV-based similarity using ROI-based analyses ^62^). This is partly because investigating such questions require a large neuroimaging dataset of parent-child dyads with rich behavioural phenotypes. Overall, the present study is the first to investigate whether, to what extent, and how brains of parent-child dyads are similar.

For both function and structure, the brain regions that contributed to similarity tended to be broadly distributed across the entire brain when we used whole-brain information (Figure 2c–2f), as in previous FC-based individual identification studies^12,67^. Although this tendency was retained when we only used the edges or regions within sub-networks, we observed slightly different contrasts between functional and structural information (Figure 3b and Table 2). Specifically, for function, the medial frontal and frontoparietal areas, which are known to be involved in networks related to higher cognitive function, revealed higher accuracies than structural information. In contrast, for structure, visual, subcortical, and cerebellum networks exhibited higher accuracies than function. This contrast is interesting because the prefrontal cortex is one of the last regions of the brain to reach maturation, exhibiting development until approximately 25 years of age, whereas subcortical regions reach maturation earlier^72^. Note that, for structural brain information, only a small number of studies using GMV are comparable to the current study^73,74^, and none of them investigated each network’s contribution in detail.

We confirmed that our results were not solely driven by structural or functional similarity alone. The neural similarity between FC and GMV exhibited a weak correlation (Supplementary Figure 6). In addition, combining FC and GMV led to the highest accuracy for many brain regions (Table 1). These results indicate that FC and GMV are distinct and contain complementary information. Previous studies reported that function and structure contain similar information, but also distinct information^68^. One study reported that information obtained from function and structure contain complementary information for the identification of siblings, but did not investigate the contributions of distinct anatomical brain locations^75^. The current results are not only consistent with those of previous studies but also demonstrated qualitative differences between function and structure by exhaustively investigating the contributions of each distinct anatomical brain location. It is noteworthy that GMV was able to obtain similar accuracy, even with short scans that are routinely acquired as initial scans in MRI protocols.

We also found that several demographic and behavioural factors had effects on the brain similarity of parent-child dyads. First, we found that age had an effect on accuracies of GMV, but not FC. It is known that both FC and GMV change through adolescence^76,77^; thus, this difference suggests that the functional and structural development of the brain are not qualitatively equal, at least from the perspective of parent-child similarity. Second, when we split children into males and females, we found age effects on the brains of female children, both for FC and GMV. This finding suggests that the developmental trajectory of the brain qualitatively differs between females and males. In addition, female children were more similar to their parents than male children, particularly at age 13. This finding is intriguing because previous studies also reported that female children are more similar to their mothers both behaviourally^78^ and neurally^62^. Third, testosterone affected brain similarity in parent-child dyads. Interestingly, children with high levels of testosterone exhibited greater similarity than other children, despite female children having greater accuracy than male children. Note that there were no significant differences in levels of testosterone between males and females both at age 11 (two-sample t-test, t(49) = −0.64, P = 0.53, Hedge’s g = −0.19) and 13 (two-sample t-test, t(44) = 0.81, P = 0.42, Hedge’s g = 0.25). It is known that levels of testosterone increase through adolescence, especially in male children^79^. Thus, different results may have been obtained if older children were tested. Fourth, questionnaire-based development scores had effects on similarity. Children with higher developmental problem scores were more similar to their parents than other children. Although this result is somewhat counterintuitive, it suggests that similarity of the brain does not merely represent behavioural maturity assessed by questionnaire. Overall, the current results provide the first detailed picture of the signature of intergenerational transmission in the brain.

Recent developments in cognitive neuroscience have made it possible to investigate individual differences, an issue that has not been deeply investigated because of the absence of adequate datasets, analytical techniques, and computational resources. The current results shed light on the importance of investigating family-level differences, in addition to individual-level differences. Families are not a neutral environment for identity development, but deeply affect individuals from adolescence, strongly influencing the development of a person’s identity^80^. Given that the current results revealed that parental brains are similar to the brains of their children, we propose that future studies should investigate the relationships between family-level behavioural indices and family-level brain information to enable more reliable predictions of children’s behaviour and development. We believe our dataset will help to extend research in this direction^64^.

The current findings indicate several potentially interesting questions for future research. The first and perhaps most important question is whether parent-child brain similarity changes across children’s development. Recent studies proposed a brain-based quantitative approach for investigating the trajectory of development^81,82^. Although the current study revealed that children at age 11 and age 13 are different, it may be valuable to investigate whether the trajectory of the parent-child brain similarity affects the various risks faced by adolescents, including psychiatric disorders and criminal behaviour, using much longer-term longitudinal datasets. Second, insights may be gained by using genetic information to confirm the extent to which genetic factors contribute to neural similarity. We examined biological parent-child relationships in the current study. Thus, it may be valuable to test whether non-biological parents and children show the same level of brain similarity. Third, we used GMV as a structural brain measurement because a number of previous studies investigated the relationship between GMV and various traits. However, the human brain also exhibits individual differences in white matter microstructure. Diffusion tensor imaging provides measures of white matter integrity in the brain, and can provide useful data, but, like GMV, produces different information to FC^83^. Indeed, previous studies have shown that thickness correlations partially reflect underlying fibre connections but contain exclusive information^84^. Future studies should use other types of structural information, such as structural connectivity obtained by diffusion tensor imaging. Fourth, because our dataset mostly consisted of mothers (81 mothers among 84 parents), it may be valuable to test whether we can also identify children’ brains from fathers. Although we confirmed that we were able to successfully conduct analyses both male and female children, future studies should investigate the effects of parents’ sex on performance.

In the present study, we sought to address a critical question in social science: whether, to what extent, and how parents and children are similar. Our analytical framework and the richness of our dataset made it possible to ask the question from the neurobiological perspective. The results revealed that parents’ and their children’s brains exhibit a high degree of similarity, and that various factors, including age, sex, hormones, and development score have effects on similarity. These results provide a comprehensive picture of the neurobiological substrates of parent-child similarity, and show the usability of our dataset for investigating the neurobiological substrates of intergenerational transmission.

## Acknowledgments

This study is the result of the projects “Science of personalized value development through adolescence: integration of brain, real-world, and life-course approaches” (16H06396, 16H06398, 16H06399) and “Adolescent Mind & Self-Regulation” (23118001, 23118002) of Grants-in-Aid for Scientific Research on Innovative Areas from the Japan Society for the Promotion of Science. We thank Hiroshi Imamizu, Ph.D., Ryu Ohata, Ph.D., Ayumu Yamashita, Ph.D., and Aurelio Cortese, Ph.D. for reviewing the draft and helpful discussions. We thank Okito Yamashita, Ph.D., and Jun-ichiro Hirayama, Ph.D. for helpful discussions. We thank Kana Inoue and Kaori Tachi for their support with data curation. We thank Giuseppe Lisi, Ph.D. for his support with code sharing. We thank Corey Horien for his support with code and data sharing.

## Author Contributions

N.O., S.A., N.Y., K.M., D.K., S. Kawakami, K.S., S. Koike, K.E., S.Y., A.N., K.K., and S.C.T. recruited participants for the study and collected their behavioural and imaging data. N.O. inspected imaging data. N.Y. provided codes for preprocessing resting state fMRI data. S. Koike, and K.K., interpreted the results and provided technical advice. S.C.T. supervised the study. Y.T. conceptualised the study, conducted the analysis, visualised the data, and wrote the manuscript. All authors reviewed the manuscript.

## Data and code availability statement

The data that support the findings of the current study may be available from the corresponding author upon reasonable request (http://value.umin.jp/data-resource.html). The code supporting the findings of this study will be published after acceptance.

## Methods

### Overview of the dataset

The Tokyo TEEN Cohort (TTC) study, which was launched in 2012, is a large-scale longitudinal general population-based survey to elucidate puberty development during adolescence, particularly the acquisition processes of self-regulation and willingness to face challenges, by focusing on the interaction between biological, psychological, and social factors^63,64^. This study was conducted as part of the population-neuroscience component of the TTC (pn-TTC) study, in which 301 early adolescents were recruited from the general population. Subjects of the pn-TTC study were subsampled from a larger subject group of the TTC study, and it was confirmed that the pn-TTC subsample was representative of the TTC study population. Written informed consent was obtained from each subject and the subject’s primary parent before participation. All protocols were approved by the research ethics committees of the Graduate School of Medicine and Faculty of Medicine at the University of Tokyo, Tokyo Metropolitan Institute of Medical Science, and the Graduate University for Advanced Studies. All research was performed in accordance with relevant guidelines/regulations. The detailed methods for subject recruitment are described elsewhere^63,64^. The dataset is publicly shared upon request (http://value.umin.jp/data-resource.html).

We excluded subjects who exhibited anomalies in fMRI or T1w images. We also excluded parents who did not have either fMRI or T1w images, and children who did not have either fMRI or T1w images at age 11 and age 13. After this screening process, 84 dyads were included in the final analysis (39 female children; 81 mothers; age = 11.59 ± 0.66 for children at age 11, 13.63 ± 0.62 for children at age 13, and 43.35 ± 0.62 for parents, mean ± s.t.d).

### MRI parameters

Subjects were instructed to lie supine on the bed of the MRI scanner. MRI scanning was performed on a Philips Achieva 3T system (Philips Medical Systems, Best, The Netherlands) using an eight-channel receiver head coil. Each subject underwent resting-state fMRI and T1-weighted (T1w) three-dimensional magnetisation-prepared rapid gradient echo (3D-MPRAGE) sequences.

Sagittal T1w images were acquired using the 3D-MPRAGE sequence with the following parameters: repetition time (TR) = 7.0 ms, echo time (TE) = 3.2 ms, minimum inversion time = 875.8 ms, flip angle = 9°, matrix = 256 × 256, field of view (FOV) = 256 mm × 240 mm × 200 mm, voxel size = 1 mm × 1 mm × 1 mm, slice thickness = 1 mm, number of slices = 200. The acquisition time was approximately 10 min 42 sec.

Resting-state fMRI images were acquired using a gradient-echo echo-planar imaging (EPI) sequence with the following parameters: TR / TE, 2500 ms / 30 ms; flip angle, 80°; matrix, 64 × 64; FOV, 212 mm × 199 mm × 159 mm; voxel size, 3.31 mm × 3.31 mm; slice thickness, 3.20 mm; slice gap, 0.8 mm. Each brain volume consisted of 40 axial slices and each functional run contained 250 image volumes preceded by four dummy volumes, resulting in a total scan time of 10 min 40 sec. Subjects were instructed to stay awake, to keep their minds as clear as possible, and to keep their eyes on a fixation point at the centre of the screen through a mirror during scanning.

### Information extraction

We used Statistical Parametric Mapping 8 (SPM8: Wellcome Department of Cognitive Neurology, http://www.fil.ion.ucl.ac.uk/spm/software/) in MATLAB (MathWorks, Natick, Massachusetts) for preprocessing and statistical analyses.

#### *Preprocessing of* structural *MRI*

T1w images were segmented into three tissue classes (grey matter [GM], white matter [WM], and cerebrospinal fluid [CSF]) using a segmentation approach implemented in SPM8. The segmented images (only GM) were then normalised into standardised Montreal Neurological Institute (MNI) space by applying a deformation field in SPM8. The GMV of each ROI was extracted and averaged within that ROI. We used a functional atlas defining 268 ROIs that cover the entire brain (functional atlas from Finn et al.^12^, which used the method developed by Shen et al.^65^) (this atlas can be downloaded from https://www.nitrc.org/frs/?group_id=51), enabling us to obtain a vector with a size of 268 for each subject. Note that, although an alternative method for inter-subject registration called Diffeomorphic Anatomical Registration Exponentiated Lie algebra (DARTEL) exists, we did not employ it because our goal was not to conduct comparisons at the group-level. Future studies should investigate whether employing another segmentation and normalisation method can improve accuracy.

#### Preprocessing of resting-state fMRI

Preprocessing of resting-state fMRI included slice-timing correction, realignment, co-registration, normalisation to MNI space, and spatial smoothing with an isotropic Gaussian kernel of 6 mm full-width at half-maximum. To avoid the effects of head motion artefacts, we calculated framewise displacement (FD). FD is defined as the mean relative displacement between two consecutive volumes for each of the six motion parameters. We conducted a “scrubbing” procedure by removing volumes with FD > 0.5 mm, along with the previous volume and two subsequent volumes, as proposed by Power et al.^66^ The average grey matter time-course for each ROI was calculated, then temporally filtered using a first-order Butterworth filter with a pass band between 0.01 Hz and 0.08 Hz. The time-course of each ROI was linearly regressed by the temporal fluctuations in white matter, cerebrospinal fluid, and the entire brain, as well as six head motion parameters. The time-course of white matter and cerebrospinal fluid were filtered using a first-order Butterworth filter with a pass band between 0.01 Hz and 0.08 Hz, and a white matter mask was eroded by one voxel to consider a partial volume effect. All parameters were determined in accord with a previous study^19^. For each subject, an FC matrix between all ROIs was then calculated by evaluating pair-wise temporal Pearson’s correlations of blood-oxygenation level dependent time courses, based only on the remaining images after the scrubbing step above. We used the same 268 ROIs that were used for GMV. Because FC matrices are symmetrical, values on only one side of the diagonal were kept, resulting in 35,778 unique edges (268 × 267/2). We then regressed the motion and total grey matter volume and mean FD out from data matrices.

#### Motion index

In addition to resting-state fMRI and structural MRI information, we performed the same analyses using motion estimates during resting-state fMRI to investigate the effects of motion artefacts^12^. We first specified 20 bins to span {0:0.05:1} to calculate discrete motion distribution vectors for each parent and child based on FD over an entire scan. These motion distribution vectors were then used in the same way as the FC or GMV vectors.

### Similarity analysis

We modified a connectome fingerprinting approach by Fin et al^12^. They used two datasets consisting of the same individual but different task sessions, called “source” and “target” dataset. They correlated the connectivity vector from one participant in the source dataset to the vectors of all participants in the target dataset and identified the maximum correlation. If the two vectors showing the strongest correlations came from the same individual, the resulting binary accuracy was 100%, whereas binary accuracy was 0% otherwise. Although these studies successfully identified brain networks that contributed to the individual identification, their method treats second-ranked and worst-ranked cases as equally failed cases, thus discarding some information that might be useful for improving the statistical power. In addition, the chance rate depends on the number of samples, thus making the interpretation difficult, especially comparing different datasets with different sample sizes. To overcome these issues, we modified the method as follows. We confirmed that the proposed method is more sensitive than conventional methods (Supplementary Figure 5) and the chance rate was always 50%, irrespective of the sample size.

For each parent-child dyad, we first calculated the similarity of their FC and/or GMV patterns based on their Pearson’s correlation. We next assessed whether the similarity of the parent-child dyad (child and their own parent) was larger than a stranger-child dyad (child and another child’s parent). We then calculated the winning rate of the similarity between parent-child dyad, denoted as “accuracy”. We repeated this procedure across all children and averaged accuracies.

Intuitively, the obtained statistics can be considered as a “pairwise classification accuracy” calculated by the following procedure:

1. Select a child randomly from all children in the sample.
2. Select two parents, including the child’s own parent and a randomly selected parent in the sample.
3. If the Pearson’s correlation coefficient between parent-child dyad is higher than that of stranger-child dyad, the result is recorded as a correct parent-child identification.
4. In contrast, if the Pearson’s correlation coefficient between parent-child dyad is smaller than that of stranger-child dyad, it is recorded as a failed identification.
5. Repeat this procedure and calculate accuracy across repetition.

By increasing the number of repetitions, this approach converges to the accuracy obtained by the main analysis. The chance rate of this approach is always 50%.

We performed 1,000-times bootstrapping to estimate the 95% confidence interval of accuracy, by randomly subsampling 90% of the subjects in each iteration. To determine whether accuracy was achieved at above-chance levels, we used 1,000-times permutation testing to generate a null distribution by randomly shuffling the parent-child mapping.

To determine the role of specific edges/regions in the performance, we quantified highly unique and highly consistent edges/regions using a differential power (DP) measure and a group consistency measure (*ϕ*) described in detail elsewhere^12^. DP provides an estimate, for each given edge/region, of the likelihood that within parent-child dyad similarity (between a parent and their parent) is higher than stranger-child similarity (between a parent and another parent’s child). Specifically, we computed the edge/region product vector (φ_*i*_) from two sets of FC/GMV vectors 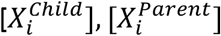,

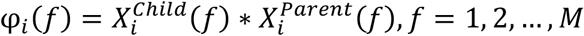

where *i* indexes dyad, *f* indexes edge/region, and *M* is the total number of edges/regions in the entire FC/GMV vector. We can calculate φ_*i*_ between vectors of a child and another child’s parent

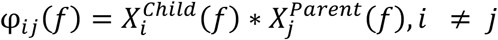

To compute the DP for all the dyads in a given dataset, we calculate an empirical probability

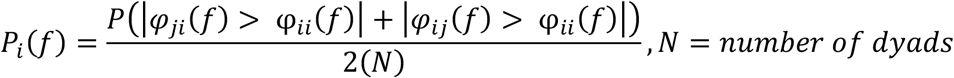

A low *P*_*i*_(*f*) indicates a more discriminative edge/region. We can finally calculate DP of an edge/region across all children in a sample:

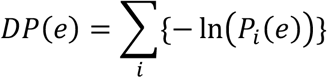

If the parent-child dyad product was higher than the stranger-child product across all children in a sample, this corresponds to a high DP value, and the edge/region is helpful.

The group consistency measure, *φ* is simply the mean of φ_*i*_ for a given edge/region across all children. Edges/regions with high *φ* values are therefore high across all pairs of children and parents, and thus are not helpful.

For the analyses in Figure 2, we used whole-brain FC or GMV. For the analyses of data shown in Figure 3, we split the whole brain into 10 sub-networks and conducted the same analyses using FC or GMV within each sub-network. The definition of sub-networks was obtained from Horien et al.^67^.

### Testosterone and Child Behavior Checklist

We investigated the effects of testosterone and Child Behavior Checklist (CBCL) scores on the brain similarity of parent-child dyads. The detailed methods for data collection are described elsewhere^63,64^. In the main analyses, we excluded dyads if either parent or child did not have a score. We also excluded children who had an extremely high testosterone measurement (more than mean + 1.5 s.t.d. Mean values of excluded children were 41.65 pg/mL [N=4] and 92.63 pg/mL [N=7] for children at age 11 and 13, respectively). After exclusion, the number of dyads was 51 for testosterone at age 11 (4.01 ± 3.63 pg/mL, mean ± s.t.d), and 46 for testosterone at age 13 (15.96 ± 17.83 pg/mL, mean ± s.t.d), 82 for CBCL at age 13 (6.55 ± 6.16, mean ± s.t.d). There was no exclusion for CBCL age at 11 (10.48 ± 9.10, mean ± s.t.d).

1. *Testosterone:* The adolescents collected their salivary samples at home early in the morning. In advance, both the adolescents and their primary parents were informed of how to collect the adolescents’ saliva using sample tubes. The adolescents tried it under the guidance of the survey staff for practice. They were instructed not to collect the saliva within a week after a tooth extraction or immediately after dental treatment to avoid contamination with blood. They were also asked not to eat food after brushing their teeth on the night before the saliva collection. They were instructed to rinse their mouth soon after getting up and to make sure they were at their normal body temperature, and not to have breakfast and not to brush teeth before the collection. Furthermore, they were asked to wait for 20 min after the rinse and then to collect 4.5 ml of their saliva by passive drool in sterilized tubes (1.5 ml/tube * 3 tubes) made of polypropylene (Nalgene^TM^ General Long-Term Storage Cryogenic Tubes, Thermo Fisher SCIENTIFIC, U.S.A.) within 60 min. Salivary samples were collected in only one day, since high correlation among morning salivary testosterone levels across days in adolescents was reported^85^. Salivary samples were kept in household refrigerator freezers, delivered frozen to our laboratory, where the weights were measured and tubes stored at minus 80 degrees C until the testosterone levels were measured. The concentration of salivary testosterone was measured once by liquid chromatography- tandem mass spectrometry (LC–MS/MS), which has become the current standard^86^. All testosterone measurements were then square-eroot transformed to better approximate a normal distribution prior to quantitative analyses.
2. *CBCL:* CBCL is a parental-report questionnaire used to screen children for behavioural problems: there are 20 competence items and 120 items on behavioural and emotional problems. The CBCL includes the following eight empirically-based syndrome scales: 1) Aggressive Behavior, 2) Anxious/Depressed, 3) Attention Problems, 4) Rule-Breaking Behavior, 5) Somatic Complaints, 6) Social Problems, 7) Thought Problems, and 8) Withdrawn/Depressed, as well as summary scores reflecting “Internalization” and “Externalization.” We used the average scores of “Internalization” and “Externalization” in the main results.

---

**Supplementary Figure 1:**
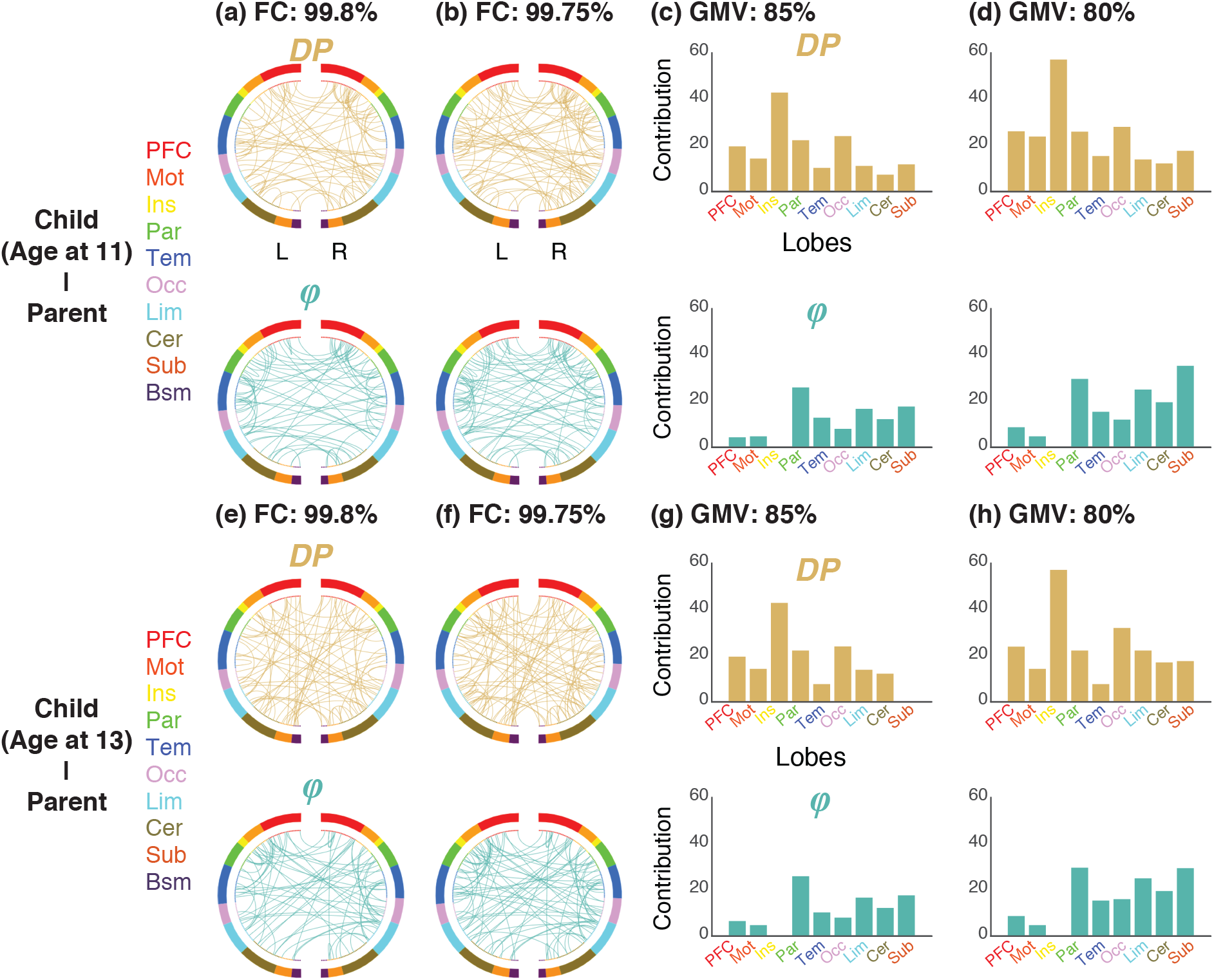
Overall patterns of edges/regions with high differential power (highly discriminative among subjects) and group consistency (highly similar among subjects) tended to be similar across different thresholds and across development (**(a)**–**(d)** child at age 11); **(e)**–**(h)** child at age 13). The figures show the results when the edges were thresholded at the 99.8th and 99.75th percentiles for FC, and regions were thresholded at the 85th, 80th percentiles for GMV. For each threshold, a circle plot (FC) and a bar graph (GMV) are shown, in which nodes are grouped according to anatomical location.

**Supplementary Figure 2:**
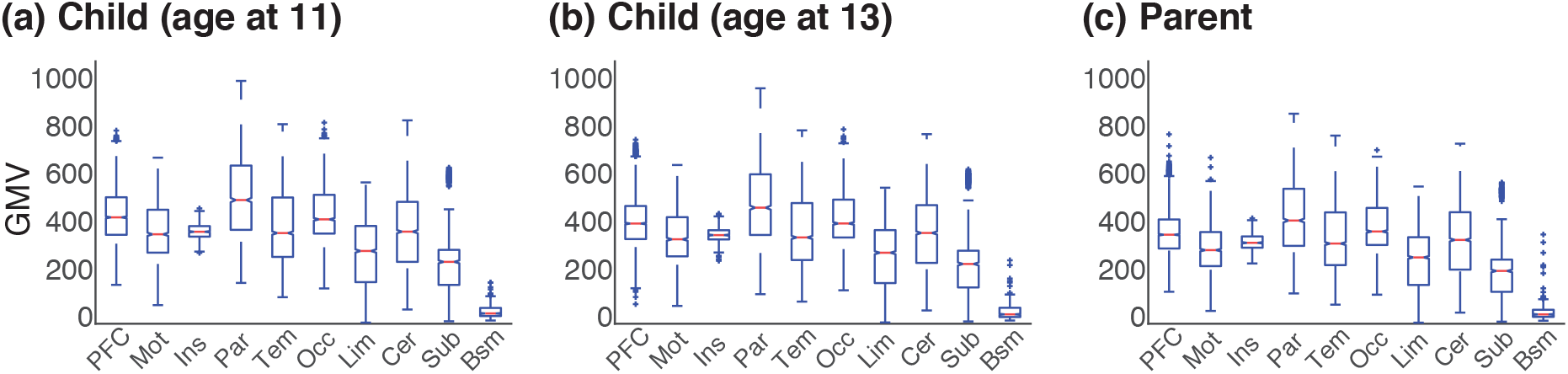
Box plots of signal strength of each lobule for GMV. The red line indicates the median. The bottom and top edges of the box indicate the 25th and 75th percentiles, respectively. The crosses denote outliers, and the whiskers extend to the most extreme data points not considered outliers. The brainstem (the right-most boxes in each figure) exhibited much lower signal strength compared with the other lobules. Means ± s.e.m are shown.

**Supplementary Figure 3:**
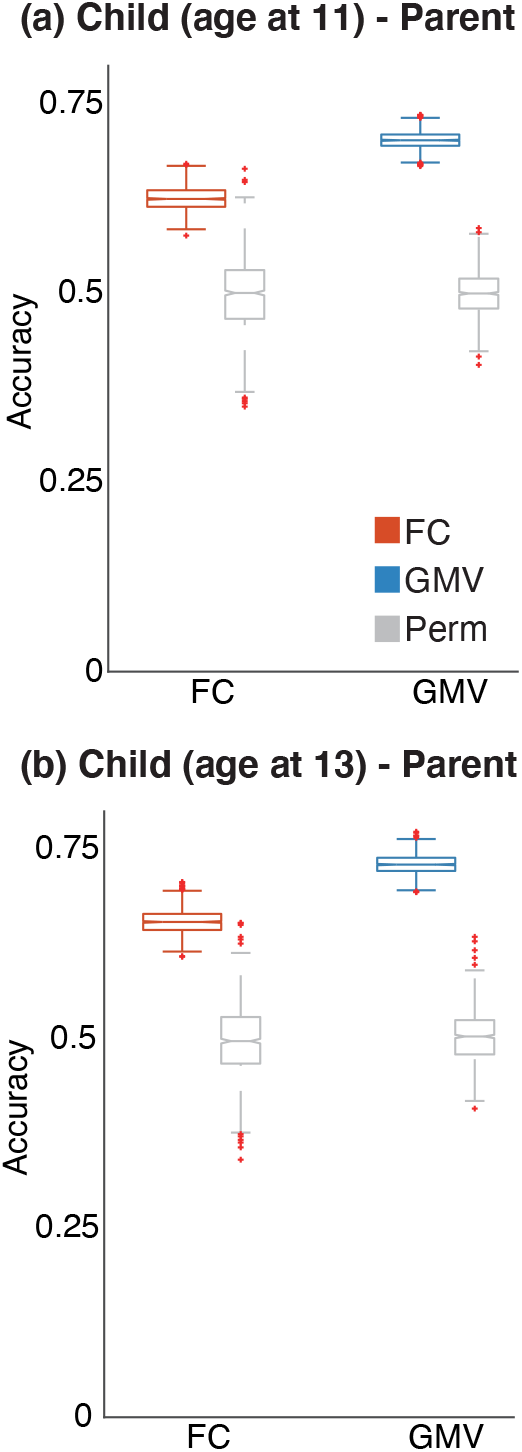
Accuracies using only low-movement subjects. We confirmed that qualitatively similar results were obtained when we excluded children whose head movements were in the top 25%, at both age 11 and age 13, resulting in the inclusion of 41.25% of the total sample.

**Supplementary Figure 4:**
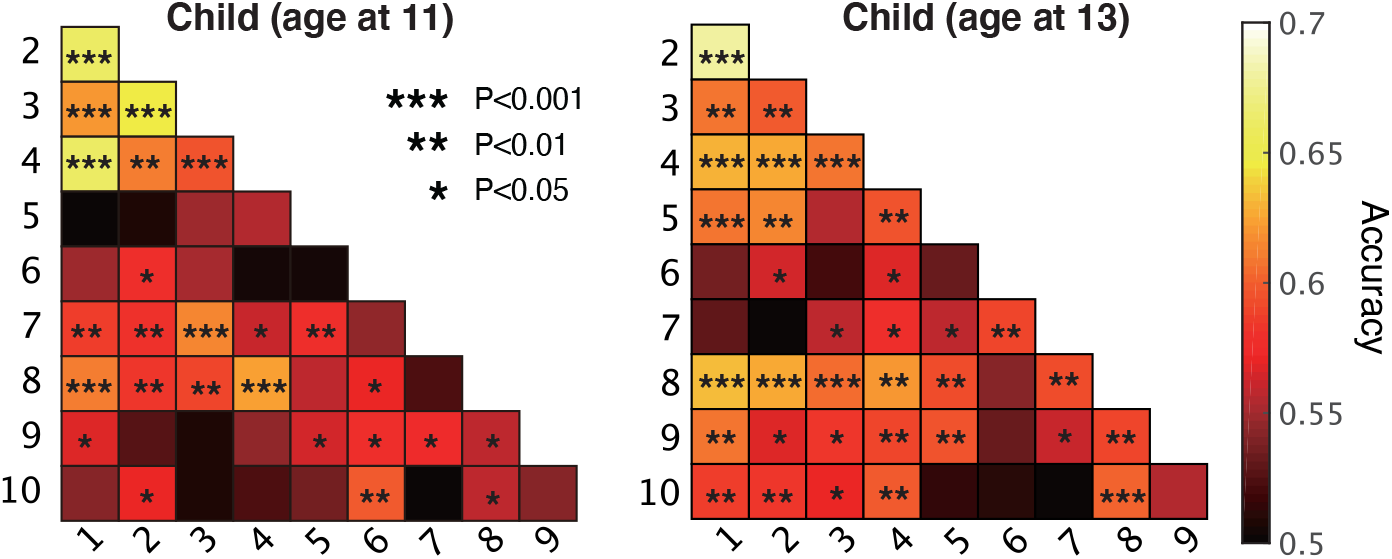
Between-network analyses using only the between-network edges. Statistical significance was assessed by comparing the distributions for each network obtained through bootstrapping, uncorrected.

**Supplementary Figure 5:**
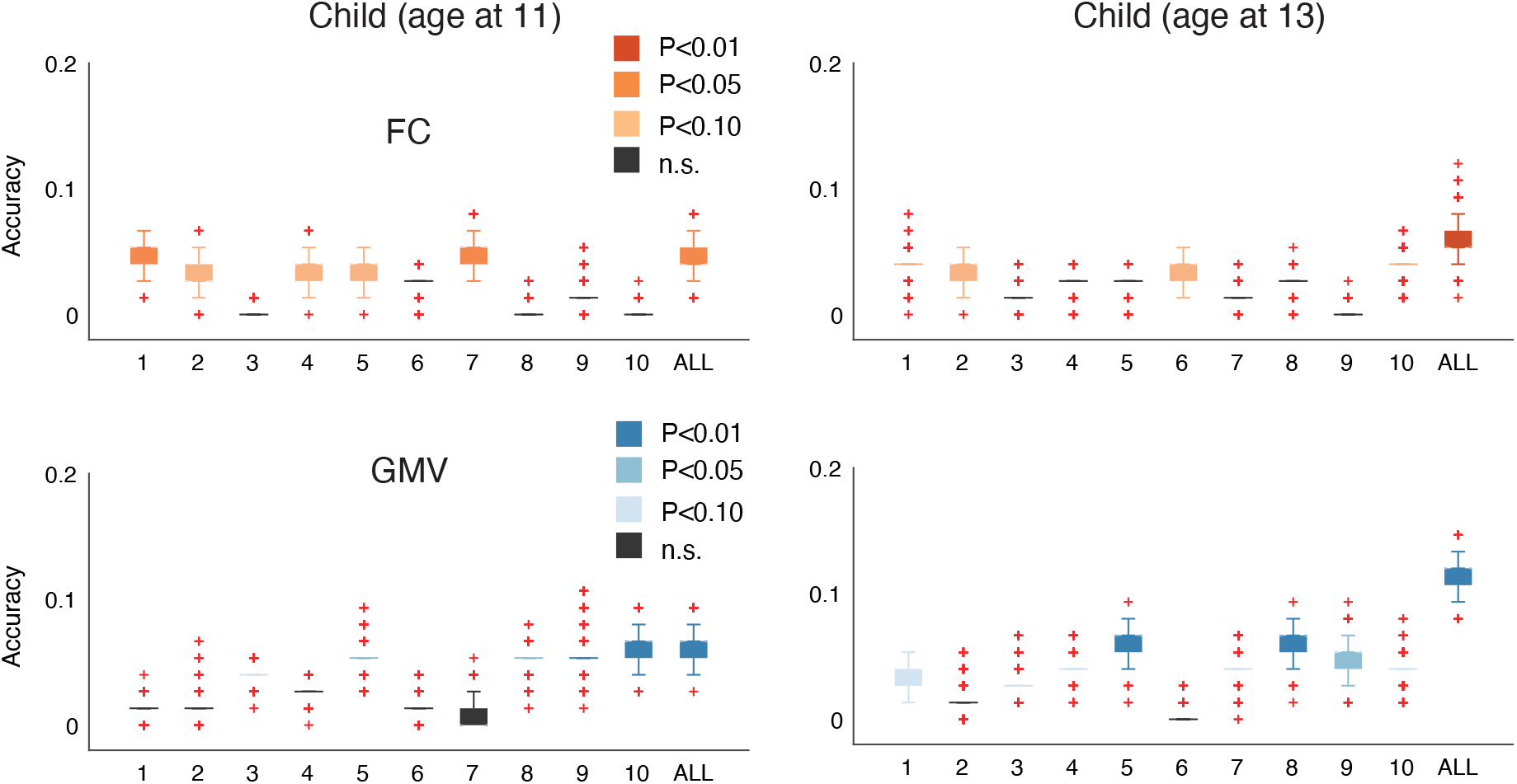
Accuracies obtained using conventional identification methods. Box plots of accuracies using FC (top row) and GMV (bottom row) for children at age 11 (left column) and 13 (right column) are shown. The bottom and top edges of the box indicate the 25th and 75th percentiles, respectively. The crosses denote outliers, and the whiskers extend to the most extreme data points not considered outliers. n.s. non-significant.

**Supplementary Figure 6:**
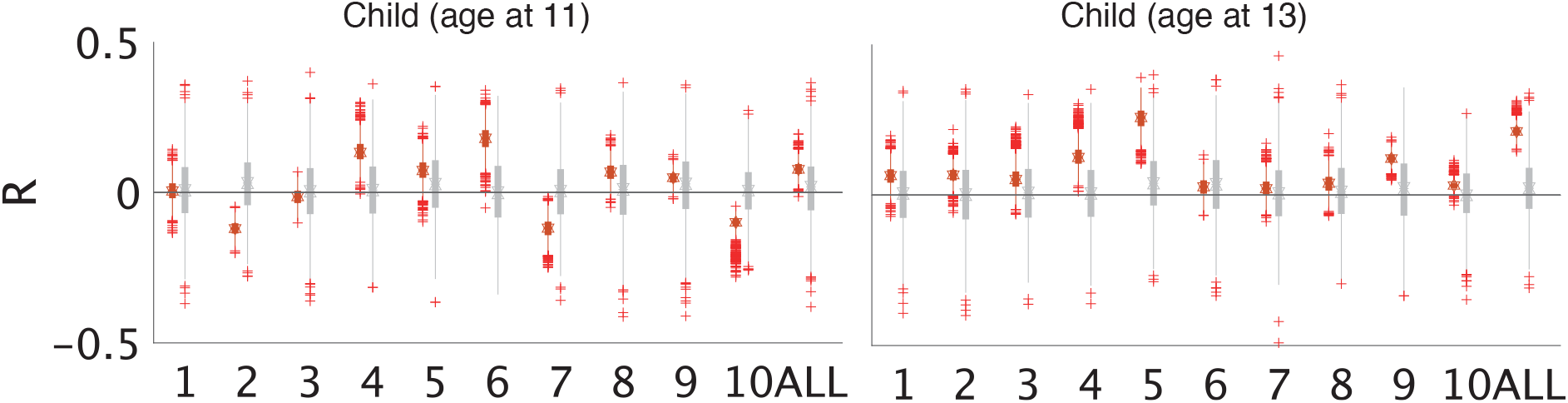
For each sub-network (1–10: sub-networks shown in Figure 3a) and whole-brain (ALL), we calculated Pearson’s correlation coefficients between parent-child brain similarities defined by FC and those defined by GMV across all parent-child dyads (red box). We found that they exhibited relatively low correlations in general, suggesting that they contained independent information. Directly to the right of these red boxes (grey box) are the results of the 1,000-times permutation testing.

**Supplementary Figure 7:**
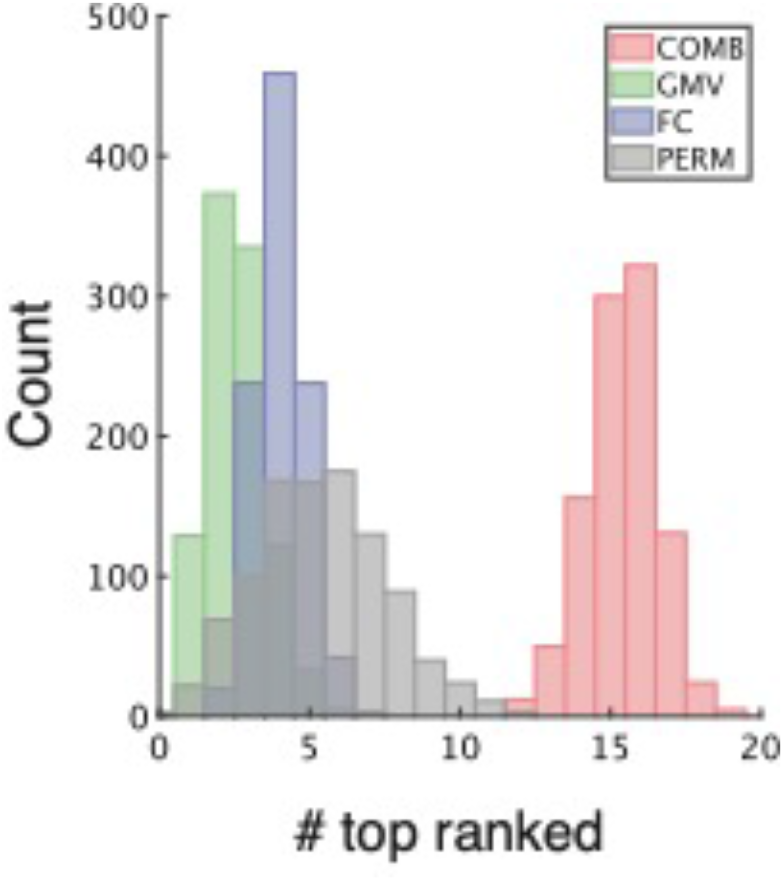
The number of top ranked networks for COMB (red bar), GMV (green bar), and FC (blue bar) estimated via bootstrapping and null distribution (grey bar) estimated via permutation are shown.

## Notes

### Competing Interest Statement

The authors have declared no competing interest.

